# MiSiC, a general deep learning-based method for the high-throughput cell segmentation of complex bacterial communities

**DOI:** 10.1101/2020.10.07.328666

**Authors:** Swapnesh Panigrahi, Dorothée Murat, Antoine Le Gall, Eugénie Martineau, Kelly Goldlust, Jean-Bernard Fiche, Sara Rombouts, Marcelo Nöllmann, Leon Espinosa, Tâm Mignot

**Author notes:** **Correspondence to:** Leon Espinosa Tâm Mignot.

## Abstract

Studies of microbial communities by live imaging require new tools for the robust identification of bacterial cells in dense and often inter-species populations, sometimes over very large scales. Here, we developed MiSiC, a general deep-learning-based segmentation method that automatically segments a wide range of spatially structured bacterial communities with very little parameter adjustment, independent of the imaging modality. Using a bacterial predator-prey interaction model, we demonstrate that MiSiC enables the analysis of interspecies interactions, resolving processes at subcellular scales and discriminating between species in millimeter size datasets. The simple implementation of MiSiC and the relatively low need in computing power make its use broadly accessible to fields interested in bacterial interactions and cell biology.

---

Bacterial biofilms and microbiomes are now under intense study due to their importance in health and environmental issues. Within these spatially-structured communities, analysis of cell-cell interactions requires powerful descriptive tools to link molecular mechanisms in single cells to cellular processes at community scales. Microscopy-based imaging methods combining multiple imaging modalities (e.g. bright-field, phase-contrast microscopy, fluorescence microscopy) directly record morphological, spatio-temporal, and intracellular molecular data in a single experiment. However, extraction of quantitative high-resolution information at high-throughput requires accurate, automatized computational tools. Methods such as MicrobeJ ^1^ and Oufti^2^ are highly performant to study single bacterial cells, but they are ill-suited to perform automated segmentation of dense bacterial communities, mostly because intensity-based segmentation is poorly applicable when the bacteria are in tight contact. The Oufti toolbox can segment single bacteria within micro-colonies, but requires extensive hand-tuning of multiple parameters limiting its robustness for high throughput, automatic data extraction^2,3^.

Machine-learning based techniques are powerful alternatives to overcome the limitations of traditional segmentation approaches. However, these techniques necessitate training which requires a large body of ground truth data often produced for a particular bacterial species and under specific imaging modalities, thus limiting the breadth of their application^3,4^. To develop a tool that can be generally applicable to studies of bacterial communities, we used a convolutional neural-network (CNN)-based segmentation method (**Mi**crobial **S**egmentation **i**n dense **C**olonies, MiSiC) that can segment densely packed bacterial cells of different species while being insensitive to the microscopy source and modality. Specifically, MiSiC is based on U-net, a CNN encoder-decoder architecture that has previously been applied for detection and counting of eukaryotic cells^5^. U-net type architectures are attractive when ground truth data is scarce, because the embedded skip-connections allow the convolutional kernels between both encoder and decoder ends to be shared^5^. This property allows fast learning from a relatively small body of labeled data. The challenge remains that the labeled data must be representative of the broadly varying experimental conditions to produce reliable outputs: in our case, different bacterial species recorded under varying imaging modalities. Therefore, we sought to develop a prediction workflow that converts an input image taken under a given imaging modality (phase contrast, fluorescence, bright field) into a binary mask for cell bodies (Figure 1). To minimize the impact of image intensity fluctuations that inevitably arise from varying imaging sources, the input images were transformed into intermediate image representations obtained from the shape and curvature (the Hessian or second-order differentiation) of the imaged objects. This strategy is possible because in rod-shaped bacteria, the characteristic dome-shaped curvatures of the poles is remarkably conserved across division cycles^6^. The curvature changes in the intensity field of an image are thus represented in a so-called Shape Index Map (SIM) derived from the eigenvalues of the Hessian of the image^7^ (see Methods section). Therefore, all microscopy images can be transformed into SIM images with intensity values ranging from −1 to 1, with −1 representing a negative dome-shape and +1 representing a positive dome-shape^7^. We then trained a U-Net to segment images of bacterial cells acquired under various experimental conditions based on SIMs.

**Figure 1.**
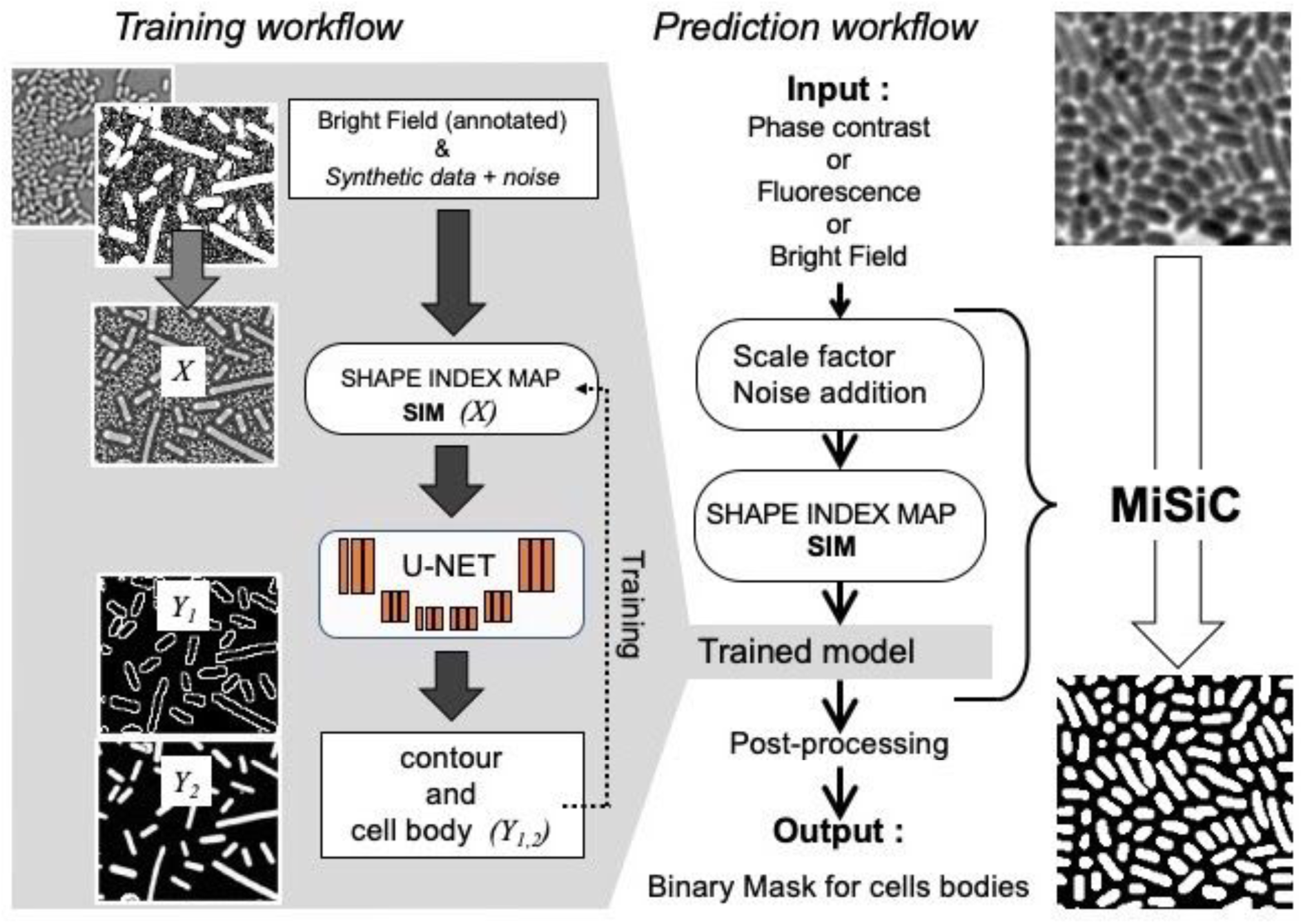
MiSiC: A U-net based bacteria segmentation tool. A set of annotated bright-field images of *Escherichia coli and Myxococcus xanthus* along with synthetic labeled data with additive Gaussian noise was used to generate a training dataset of input images, X, consisting of Shape Index Map of intensity images (at three scales) and segmented images, Y, consisting of contours (Y1) and cell body (Y2). A CNN with U-net architecture was trained to segment the Shape Index Maps into cell body and contour of bacterial cells. Prediction using MiSiC requires that the width of the bacteria in the input image be contained in 9-11 pixels, which is easily obtained by rescaling the input image based on the average width of the bacteria under consideration. Gaussian noise may be added to the input image to reduce false positives (Methods).

A schematic of the training strategy is shown in Figure 1 and detailed in the Methods section. Specifically, the U-Net was trained to segment cells by learning shapes of individual bacteria and patterns emerging from the tight contact between cells. Accordingly, we curated a hand-segmented dataset of 128 bright-field images of two rod-shaped bacterial species, *Escherichia coli* and *Myxococcus xanthus*. We further enriched the dataset with synthetic data obtained with a simple model for rod-shaped bacteria with a 0.5 µm width corresponding to 8-10 pixels in the image. The ground truth data has two classes: one with the mask of bacteria and the other with the contour of the detected cell (Figure 1). This allows other algorithms like watershed, conditional random fields, or snake segmentation to be used for post-processing to separate bacteria in cases where there is not enough edge information for proper separation of tightly connected bacterial cells. Prior to segmentation, two parameters must be adjusted to generate a SIM image from an input image: (i) Due to the scale of the training data-set, obtaining a satisfactory segmentation with the trained network requires the input image to be scaled so that the average width of the bacterial cell is contained in 10 pixels. (ii), The scaling often smoothens the original image, which in turns smoothens the corresponding SIM. This is potentially problematic because the U-Net distinguishes smooth curvatures with well-defined boundaries and noise reduction can lead to increased false positive segmentation in the scaled images. We solved this problem by adding synthetic noise to the scaled images. Thus, the MiSiC workflow takes raw input images of any imaging modality, scales them and adds noise to generate SIMs that are then segmented with the above described U-Net (Figure 1).

The SIM representation effectively allows MiSiC to efficiently segment images of bacteria in dense colonies, either from phase contrast, fluorescence or brightfield modalities (Figure 2a, Figure S1a for a statistical analysis). MiSiC works equally well with noisy backgrounds often encountered with low exposure conditions required for fluorescence time-lapse imaging (Figure S1b and comparison with the performance of another available software^3^). MiSiC can also readily segment bacterial species of distinct shapes such as *Pseudomonas aeruginosa*, *Caulobacter crescentus* and filamentous *Bacillus subtilis* captured using various microscopy modalities from different laboratories (Figure 2b, Table S1). As MiSiC was trained using rod-shaped bacteria, we expected the quality of the segmentation to decay as bacterial shapes deviate from the straight rod. To quantitatively evaluate this deviation we compared manual-segmentation (Figure 2c, Methods) to MiSiC-based segmentations of bacterial species with various shapes, classical rod shapes (*E. coli* and *B. subtilis*), curved “crescent” shapes (*Caulobacter crescentus* and *Desulfovibrio vulgaris*) and non-rod shape filament-forming bacteria (*Anabaena* sp). MiSiC was able to segment a large variety of bacterial species in absence of any parameter tuning. As expected, cells with non-rod shapes *(ie Anabaena* forming septated filaments) were less robustly segmented by MiSiC (See Discussion).

**Figure 2.**
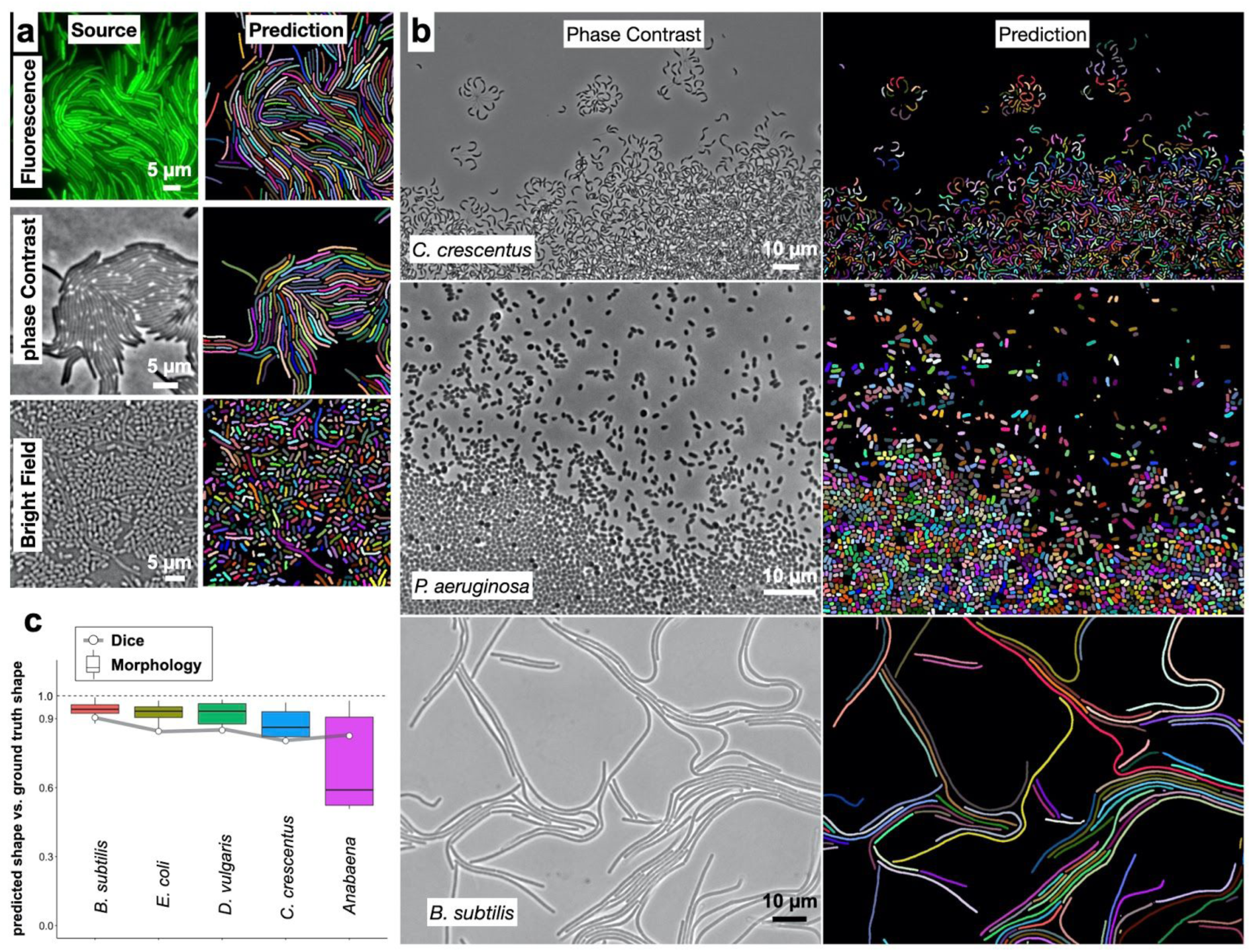
MiSiC segmentation for multiple imaging modalities and bacterial species of different shapes. a) MiSiC mask prediction of bacterial colonies captured in phase contrast, fluorescence and bright field modes. Left panels: source images, Right panels: MiSiC mask predictions. b) MiSiC mask prediction of three different bacterial species of distinct shapes: top panel: *C. crescentus* (curved), middle panel: *P. aeruginosa* (rounded small rod) and bottom panel: *B. subtilis* (filamentous). Left panels: source images, Right panels: MiSiC mask predictions. c) Accuracy assessment of the predicted shapes by MiSiC against manual ground truth segmentation. Five species with different morphologies were tested. Two parameters were calculated: the Dice similarity coefficient^8^ and a morphological index meaning geometrical features normalized to 1 for perfect identity (see Methods). As the morphology of the target species deviates from the rod shape, the variance of the morphological prediction increases and the similarity slightly decreases from 0.90 to 0.80 (Images in Figure S2)

Encouraged by these results, we tested whether MiSiC could be further used to study bacterial multicellular organization and inter-species interactions accurately at very large scales. As a model system we used *Myxococcus xanthus*, a delta proteobacterium living in soil, that predates collectively in a process whereby thousands of cells move together to invade and kill prey colonies^9^. In the laboratory, spotting a *Myxococcus* colony next to a prey colony (here *E. coli*) results in invasion and complete digestion of the prey cells in 48 H (Figure 3a). To capture predator-prey interactions at single cell resolution, we set up a predation assay where *Myxococcus* and *E. coli* interact on a 1 cm^2^ agar surface directly on a microscope slide (Figure S3a). Under these conditions, the entire invasion process occurs over a single prey cell layer allowing identification of single predator and prey cells at any given stage. This area is nevertheless quite large, and to record it with cell-level resolution, we implemented a multi-modal imaging technique termed ’Bacto-Hubble’ (in reference to the Hubble telescope and its use for the reconstruction of large scale images of the galaxies) that scans the entire bacterial community with a 100X microscope objective and reconstructs a single image by near neighbour end-joining of multiple tiles of 80 nm/pixel resolution images (Figure S3b). Application of this method requires addressing practical considerations that are detailed in the methods section. Bacto-Hubble images (phase contrast and multi-channel fluorescence) thus capture cellular processes in native community environment. We next tested whether MiSiC addresses the computational challenges posed by the analysis of such complex (dense population and mixed species) and large size data sets.

**Figure 3.**
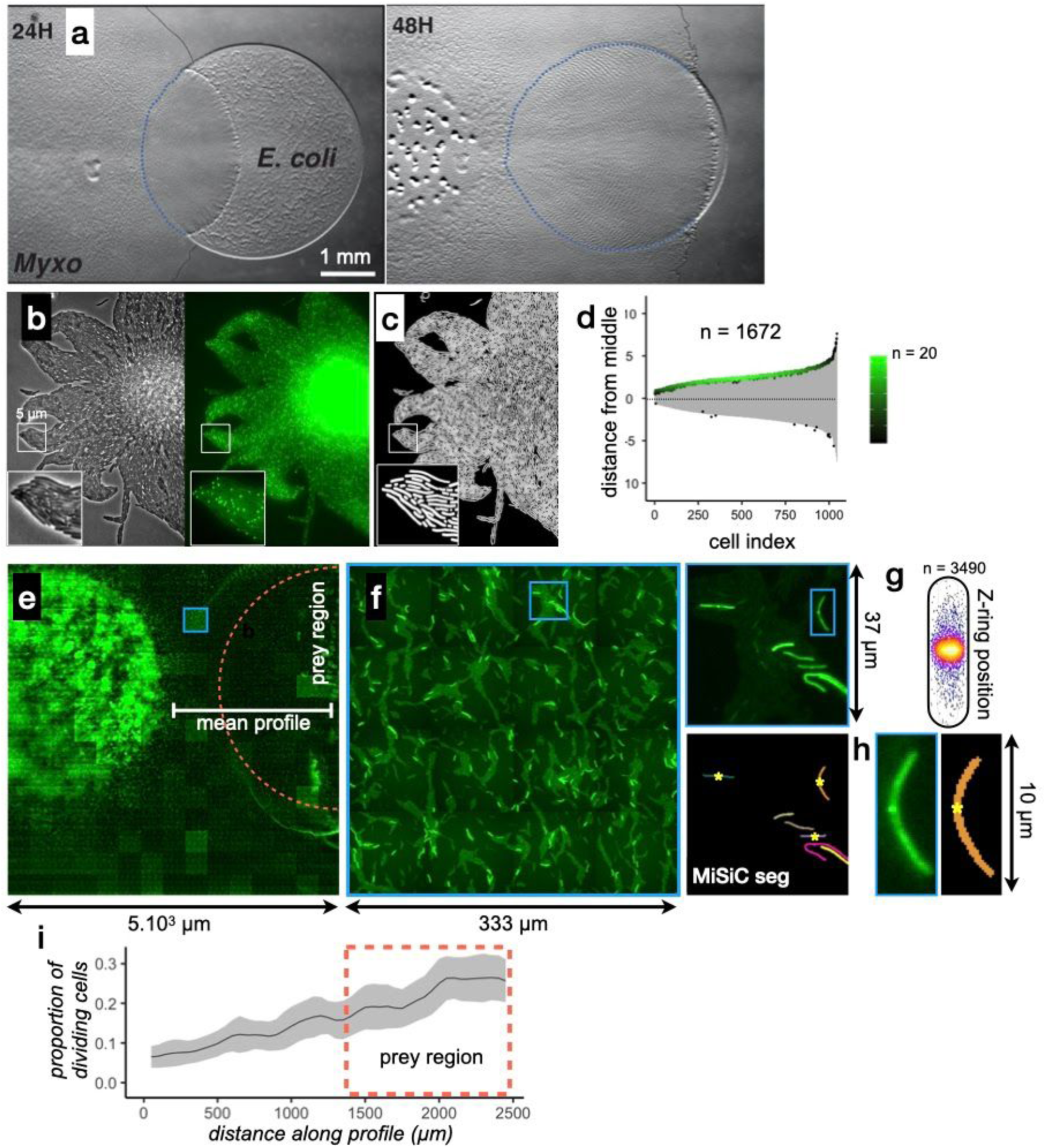
MiSiC can be applied to the study of cellular processes at the mesoscale. **a) The *Myxococcus xanthus* predation cycle on a Petri dish over the course of 96 hours.** When plated next to an *Escherichia coli* colony, *Myxococcus xanthus* uses collective movement to invade, consume and develop over the former prey colony. At 48H, spore-filled fruiting bodies are observed forming in the nutrient depleted area but not in the former prey area where the *Myxococcus* cells are actively growing. Scale bar = 0.5 mm. **(b-d) MiSiC can segment dense bacterial swarms.** b) An *M. xanthus* swarm expressing SgmX-GFP, observed at colony edges and captured under phase contrast, fluorescence and corresponding magnified images. c) MiSiC prediction mask obtained on the phase contrast image shown in b d) Demograph representation of the segmented cells and corresponding localization of the SgmX-GFP fluorescent clusters. In this representation the cells are aligned by order of length (grey area and the position of the clusters is positioned with respect to the position of the cell middle set to 0. The color of clusters reflects the histogram of the cell length distribution (bins = 0.05 µm, maximum = 20 cells for [3.9 - 3.95] µm). **(e-i) Mapping of *M. xanthus* cell division in the *M. xanthus*-*E. coli* community.** (e-f) Bacto-Hubble image of a predatory field containing FtsZ-NG labeled *Myxococcus xanthus* cells and unlabeled *Escherichia coli* prey cells. The composite image results from the assembly of 15×15 Tile images. The dotted circle marks the limits of the original prey colony. The white line (mean profile) indicates the axis used for the analysis shown in (i). (f), a single image tile showing a representative density of fluorescent cells. (g) Detection of dividing cells. FtsZ-NG fluorescent clusters are detected at midcell. The FtsZ clusters can be detected as fluorescence intensity maxima. Shown is a projection of the position of fluorescence intensity on a mean cell contour for a subset of n = cells (representing 1/7 of the total detected cells with a cluster), revealing that as expected, the clusters form at mid-cell. The blue square marks the cell shown as an example in (h). (h) Counting dividing cells. The example shows segmentation of the field shown in (h). The position of Z-ring foci detected as fluorescent maxima was linked to all fluorescent cells segmented in the MiSiC mask. (i) *Myxococcus* cells divide in the prey colony. The spatial density of dividing cells and the total fluorescent cells (Methods) and the proportion of dividing cells (density of dividing cells/density of total cells, methods) were determined all across the prey area shown in (e) (dotted circle). The mean ratio and standard deviation are plotted along a spatial axis (distance along profile) corresponding to areas outside and inside (dotted rectangle) of the prey area (mean profile, white segment in (e)).

First, we tested the capacity of MiSiC to segment closely-packed swarms of *Myxococcus xanthus* cells captured in a single image tile. To test the fidelity of segmentations in these conditions, we imaged a swarm composed of cells expressing SgmX-sfGFP, a motility protein that localizes to the cell pole^10^ in both fluorescence and phase contrast modalities (Figure 3b). Phase contrast images were used to obtain a MiSiC segmentation mask (Fig. 3c). Subsequently, the mask was filtered using MicrobeJ^1^ to remove objects that do not correspond to cells (Methods, less than 1.4 %, n=1695). Next, we calculated the localization pattern of SgmX-GFP foci with respect to the long axis of each segmented cell (Figure 3d). As expected, SgmX-GFP loci localized to a cell pole, consistent with most *Myxococcus* cells in swarms being properly segmented by MiSiC.

Second, to show that MiSiC can be used to quantitatively study cellular processes in entire Bacto-Hubble images, we mapped a *Myxococcus* cellular process directly during prey invasion. Cell division is expected to occur mostly in prey-areas in absence of any other source of nutrients. Like all rod-shaped bacteria, dividing *Myxococcus* cells assemble a polymeric FtsZ bacterial tubulin ring to initiate cell division^11^. When it is fused to fluorescent proteins, the FtsZ ring is observed as a dot at mid-cell^12^, which can be used as a proxy to determine which cells enter division. Thus, we first engineered *M. xanthus* cells expressing FtsZ fused to Neon-Green (NG, Methods^6^) and mixed *Myxococcu*s FtsZ-NG+ (5 %) with non-labeled cells (95 %) in the presence of an *Escherichia coli* prey cell colony. A fluorescence Bacto-Hubble image spanning ~5 mm^2^ (representing 225 tiles of 500×500 pixels images) of the community during the invasion phase (Figure 3e) was then captured and segmented tile-by-tile using MiSiC (Figure 3e-h). Cells with mid-cell FtsZ-NG fluorescence clusters were clearly observed suggesting that cell division is ongoing (Figure 3g). Dividing cells were therefore counted across the entire image (Figure 3h, Methods) to determine where they localize spatially within the community. Figure 3i, shows that cell division is markedly increased in the prey area, demonstrating directly that *Myxococcus* grows during prey invasion. Thus, MiSiC is appropriate for the automated detection of cellular processes (detected at subcellular resolution) at community scales.

Third, we explored the ability of MiSiC to segment and classify multiple bacterial species intermingling and interacting in space (semantic segmentation); here *Myxococcus* cell groups invading the tightly-knitted *E. coli* prey colony. To segment each bacterial species directly from unlabeled phase contrast-images, new training datasets were produced and used to retrain the U-NET. These labeled datasets were obtained by imaging GFP-labeled *Myxococcus* and mCherry-labeled *E. coli* (see Methods). Images were captured for each channel (GFP, mCherry) and segmented separately with MiSiC to obtain masks for each species (Figure 4a). These masks were used to retrain the U-NET, which was tested by acquiring a single phase-contrast Bacto-hubble image and analysing it to classify *Myxococcus* and *E. coli* cells directly in absence of any fluorescence labeling (Figure 4a-4b) We could thus discriminate cells belonging to each species in the predation area (Figure 4b) for a total detection of ~402 000 *Myxococcus* and ~630 000 *E. coli* cells in the entire image. To test the accuracy of the classification procedure, we tested whether the distribution of shape descriptors, such as the extent (E=area/bounding box area), solidity (S=area/convex area) and minor axis length, matched the distribution of these descriptors obtained from images of each single bacterial species (Figure 4c). The observed distributions were indeed consistent with an efficient separation of species in the mixed community.

**Figure 4.**
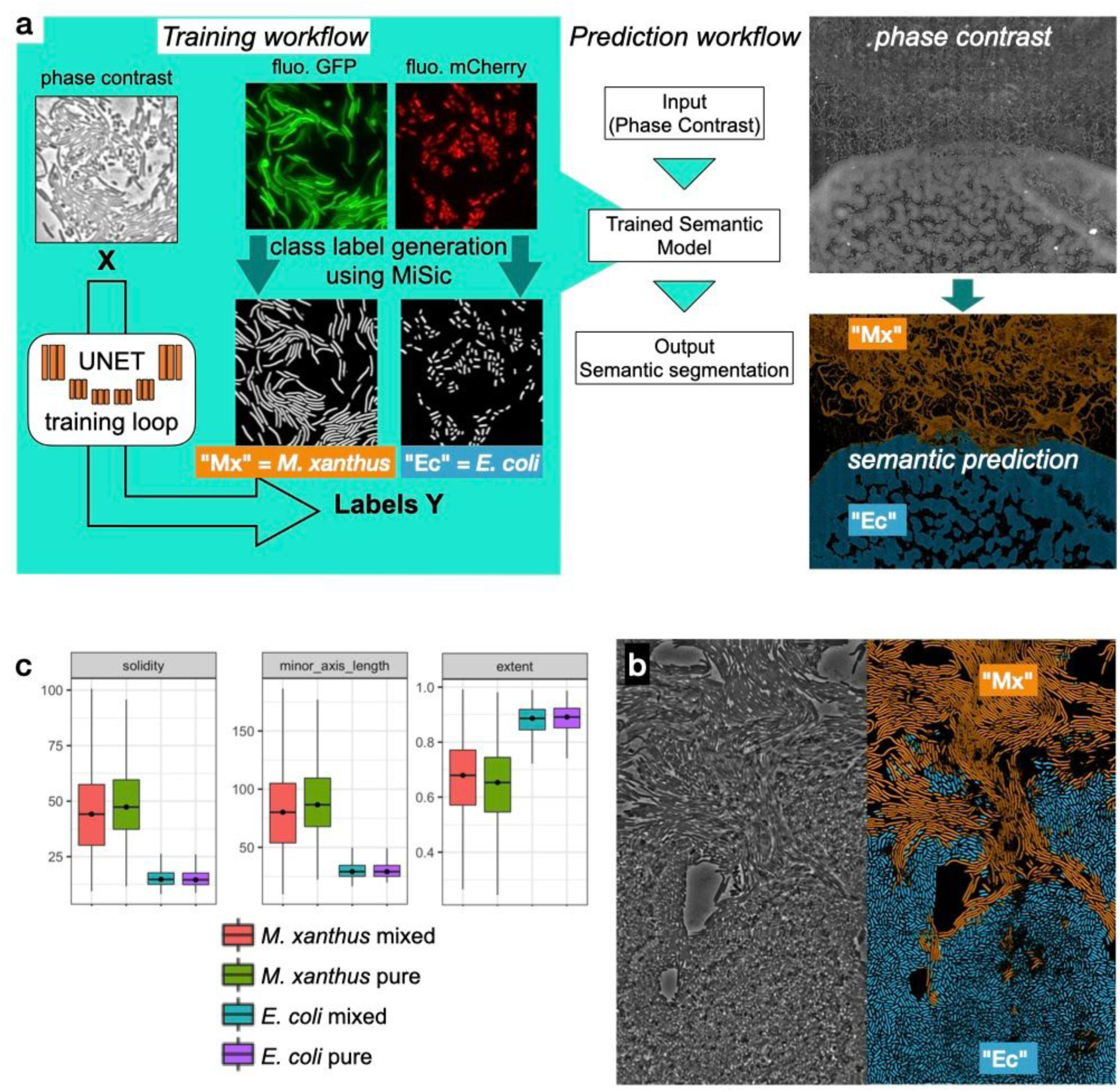
Semantic segmentation of *M. xanthus* and *E. coli* from single phase contrast images. **a) Semantic classification network.** A U-Net was trained to discriminate *M. xanthus* cells from *E. coli* cells. The training dataset consisted of GFP^+^ *M. xanthus* cells mixed with *mCherry*^+^ *E. coli* cells, which were imaged in distinct fluorescent channels (GFP and mCherry) and segmented using MiSiC to produce ground truth data for each species. The network uses unlabeled phase contrast images as input (X) and produces one output for each labeled species (Y). The prediction workflow in the right shows semantic segmentation of a Bacto-Hubble image of *M. xanthus* cells invading an *E. coli* colony after 24 hours. The composite image corresponds to 20×42 image tiles captured by phase contrast and segmented tile-by-tile to produce the resulting classification. **b-c) Direct semantic segmentation of *M. xanthus* interacting with *E. coli*.** b) Left panel: zoom of the areas where *M. xanthus* cells actively penetrate the *E. coli* colony. Left panel: Phase Contrast. Right panel: predicted mask for each species: *M. xanthus* in orange and *E. coli* in blue. The high magnification detail reveals the ability of the semantic model to label objects with low contrast in the grayscale source image. c) Morphological analyses of the classified cell population and comparison with the ground truth data. Morphological parameters (Extent, Solidity and minor axis length) were determined for the cells predicted in the *M. xanthus* (Mx mixed) and *E. coli* masks (*E. coli* mixed) in the context of a mixed colony and compared to the same parameters obtained from MiSiC segmented from images of pure cultures (Mx/E.coli pure).

## Discussion

In this article we have presented MiSiC, a deep-learning based bacteria segmentation tool capable of segmenting bacteria in dense colonies imaged through different imaging modalities. The main novelty of our method is the use of a shape index map (SIM) as a preprocessing step before network training and segmentation. The SIM depends on the Hessian of the image, thus preserving the shape of bacterial masks rather than the raw intensity values, which vary as a function of microscopy modalities. Since bacteria have simple shapes the SIM and corresponding annotations can be synthetically produced, greatly facilitating the generation of a training dataset. This strategy only requires two adjustment parameters (scaling and noise addition) and it makes segmentation agnostic to imaging modality, and adapted to different bacterial species with different morphologies, provided that they do not deviate too largely from rod shapes. For cocci, oval or rugby-ball bacteria, it should not be too challenging to further adapt MiSiC for their segmentation because their shape will also be captured in the SIM and specific synthetic training data sets could be designed. In general, the use of SIMs rather than image intensity is a promising lead for any deep learning approach to cell segmentation that relies on shape, which could also solve modality issues for eukaryotic cell and cellular organelle segmentation.

MiSiC is appropriate for the automated analysis of complex images, such as fluorescence (Bacto-Hubble) images tiles. In our hands, FtsZ-NG-expressing cells could not be properly segmented across tiles with intensity-based methods. In fact, variations in fluorescence intensities associated with multiple image captures required parameter adjustment for each tile, making the task overly complex. These problems are solved in MiSiC because it only uses a small number of parameters and it is relatively robust to noise. Importantly, tile-by-tile segmentation in MiSiC also allows extraction of high-resolution information from large size data sets with reasonable calculation power. Last, MiSiC can be combined with existing microbial cell analysis packages such as Oufti and Microbe J, which contain sophisticated tracking and analytical procedures for in-depth exploration of cellular processes at subdiffraction resolution^1,2^.

Finally, we show that MiSiC can be further implemented for the semantic classification of bacterial cell types directly from phase contrast images. While the network developed herein works specifically for *M. xanthus* and *E. coli* discrimination, the approach can be easily extended to segment and classify any number of bacterial species provided that a ground truth dataset (ie fluorescence labeling) is available to train a U-Net with ground truth generated inMiSiC. At a time where tremendous efforts are injected to reconstruct micro communities in synthetic contexts for mechanistic studies^13^, we foresee that deep learning based approaches such as MiSiC will profoundly impact microbiome research of health and environmental significance.

## Methods

### Bacterial strains and predation assays

The complete list of the strains used for the study is compiled in Table S1. For predation assays, cells of *Myxococcus xanthus* (DZ2, Table S1) were grown overnight in 100 mL flasks in 10 to 20 mL of CYE^14^ media without antibiotics at 32°C with shaking. In parallel a colony of *Escherichia coli* (MG1655, Table S1) was grown in 5 mL LB medium in a glass tube at 37°C with shaking. The next day, OD_600 nm_ were measured and cultures of both strains were washed twice at 5.000 rpm in CF^14^ minimal media to discard CYE and LB traces. After the washes, the density of the cultures was brought to 5 OD units in CF media. Pads of CF agar 0.5% were poured in precast frames (in situ GeneFrame, 65 µL, ABGene, AB-0577) that were mounted on glass slides and briefly dried. 1 µL of both *Myxococcus xanthus* and prey cell suspensions were spotted as close as possible to one other on the pad making sure that they would not merge. Glass slides were kept in a sealed humid dish for 6, 24, 48 or 72 hours at 32 degrees. 30 minutes before observation, the agar pad around the colonies was cut out and discarded and the pad was sealed with a cover slip and observed by microscopy.

*Bacillus subtilis* strains were grown in LB medium at 37°C until they reached an OD_600 nm_ of 0.6, were transferred into wells and covered in a low melting LB-based agarose suspension (2%) before observation. *Pseudomonas aeruginosa* was grown in LB medium and cells were observed at an OD_600 nm_ of 0.5. *Caulobacter crescentus* was cultured in PYE on the benchtop without shaking for 3-4 days. Cells were imaged from disrupted fragments of a pellicle biofilm and were transferred from the air-liquid interface to glass slides for imaging^15^. *Anabaena nostoc* was grown in BG11 medium at 30°C with illumination (40 µE m-2s-1). Finally, *Desulfovibrio vulgaris* cells were grown until mid-exponential phase (OD_600 nm_ of approximately 0.4 to 0.5) in LS4D medium supplemented with 1 g/L of yeast extract (LS4D-YE) at 33°C in an anaerobic chamber (COY Laboratory Products) filled with a 10% H_2_-90% N_2_ mixed-gas atmosphere. Cultures (200 µL) were centrifuged, and the pellet was resuspended in 100 µl of 10 mM Tris-HCl (pH 7.6), 8 mM MgSO_4_ and 1 mM KH_2_PO_4_ buffer (TPM buffer). The cells were placed between a coverslip and an agar pad containing 2% of agarose.

### Molecular Biology and strain construction

To follow cell cycle progression in single cells of *Myxococcus xanthus*, a merodiploid strain of DZ2 expressing both native FtsZ and the fusion protein FtsZ-neonGreen (FtsZ-NG) was built (DM14, Table S1). To do so, the coding sequence of DZ2 *ftsZ* gene (MXAN_5597) along with its predicted promoter sequence was amplified by PCR with primers oDM1 and oDM2 (Table S2) and cloned in the non-replicative plasmid pKA32^12^ allowing for its site-specific integration at the DZ2 *attmx8* phage attachment site on the *M. xanthus* chromosome. The coding sequence of the neonGreen protein was amplified from a plasmid^16^ using primers oDM16 and oDM17 (Table S2) allowing the in frame addition of neonGreen at the C-terminus of the FtsZ protein. When grown in CYE rich medium, DM14 did not present any significant defect in growth rate or cell shape. DM14 cell size is not significantly different from that of the isogenic wild type DZ2 strain. DM14 cells were spotted on thin CF agar pads to follow FtsZ localization in axenic cultures and allowed us to confirm that cell cycle progression was accompanied with the relocalization of Ftsz-nG from being diffuse in the cytoplasm to forming a discrete fluorescent focus at mid-cell before cell septation as previously described^12^.

To generate Dataset 2 (see below), strains of *E. coli* (EC500, Table S1) and *M. xanthus* (DM31, Table S1) expressing soluble versions of mCherry and sfGFP fluorophores respectively were used. To generate DM31, a plasmid allowing for the constitutive expression of sfGFP was built (pDM14). To obtain a high and constitutive expression of sfGFP in *M. xanthus*, we sought for the closest homolog of the constitutively expressed *E. coli* EF-TU (Translation elongation factor) in *M. xanthus* genome which is MXAN_3068. The 1000-bp region upstream of MXAN_3068 (*p3068*) was amplified by PCR using oDM53 and oDM54 and cloned upstream the coding sequence of sfGFP (amplified using primers oDM61 and oDM62) in a pSWU19 plasmid. The transcriptional fusion was then integrated on DZ2 chromosome at the *attmx8* site through transformation. DM31 cells display a constitutive bright diffuse fluorescent signal in our growth conditions.

### Microscopy and Image acquisition

All microscopy images were acquired with an inverted optical microscope (Nikon TiE) and a 100x NA=1.45 Phase Contrast objective. Camera used was Orca-EM CCD 1000×1000 camera mostly set with binning 2×2. Acquisition software was Nikon NIS-Elements with specific module JOBS. Fluorescence acquisition used a diode based excitation device (Spectra-X Lumencore).

The Bacto-Hubble image is a composite image of rasters of the entire area and requires that the scanning speed must be sufficiently fast to avoid image shifts due to ongoing cell dynamics. To minimize the shift in focus from tile to tile we used Nikon perfect focus system (PFS) equipped with servo-control of the focus with an infrared LED. This is especially challenging because continuous focus alignment of the microscope slows down the acquisition times dramatically. To obtain a satisfactory compromise allowing both fast scanning and correct focusing we:

i. reduced the number of dynamic elements on the microscope set up: we replaced shutters by Light Emitting Diodes (LEDs, Spectra-X for fluorescence source and a white diode for transmitted light) which could be switched with a high frequency rate (100 kHz). In addition, a double band dichroic mirror for the fluorescent cube was used to avoid switching the filters’ turret for each snapshot.
ii. used an EM-CCD camera set to a 2×2 binning mode to reduce the size of images (500×500 pixels at 0.16 µm/px) and acquisition time, and iii) sped up the vertical movements by means of a piezoelectric stage. In its largest scanning mode, Bacto-Hubble thus captures 80×40 raster images covering a total surface of 20 mm2 (containing up to 0.8 billion pixels, an acquisition-time up to 4 hours), enabling a continuous magnification display from eye visible structures to single-cells. Individual tiles for Bacto-Hubble images were acquired with the scan large field capabilities of NIS-Elements software. A key point for Bacto-Hubble large images was the quality of the sample slide mounting. The samples were placed on a glass slide with a thin double-sided sticky frame (in situ GeneFrame, 65 µL, ABGene, AB-0577). An agar pad was poured inside the frame and a microliter of cells was placed on it. The chamber was closed with a glass cover slide. This assembly allows very good flatness and rigidity.

### U-NET as Base architecture

We implement a U-net inspired in encoder-decoder architecture with skip-connections and use it as a base network for segmentation tasks. This architecture is now widely used for segmentation tasks and has many advantages that are discussed in previous articles ^5,17^. The original U-net architecture^5^ was modified to include relu activation for all layers except for the output layer where sigmoid activation was used. The general U-net was implemented in Python programming language using tensorflow (https://www.tensorflow.org/tutorials/images/segmentation), where the number of input channels (say, n) and output classes (say, m) could be varied as required by different models. The number of encoder layers were fixed to 4 with filter lengths [32,64,128,256] for the encoder side. The loss function was also modified from the original implementation. A combination binary cross entropy and the Jaccard index^18^ was used as the loss function with Adams optimizer (learning rate = 0.001) for the minimization.

For brevity, we denote the network as a mapping between input X and output y as,, where X is a set of images with n channels and y is the output with m classes. Thus given a training data set of X_train_ with sizes (N × S × S × n) and multiclass images y_train_ (of size N × S × S × m), the generalized implementation of U-NET learns to predict the segmented image from unknown images X.

### Dataset 1

This training dataset consists of three parts: (a) 263 cases of Bright-field images of *Escherichia coli* and *Myxococcus xanthus* with segmented masks, (b) 600 cases of synthetic data (c) 200 cases of synthetically generated null cases with random gaussian noise and circular objects and (c) 200 null cases, that contain background images without bacteria taken from bright-field, fluorescence and phase contrast data. The synthetic data is generated with a simple model for rod-shaped bacteria with a width ranging from 8-10 pixels. An overlap threshold of 2% was used to obtain dense cell population. The binary mask created is then smoothed with a Gaussian and Gaussian noise is added to emulate noise in real images. The ground truth in this training dataset (denoted as [X’, y]) has two classes: one with the mask of bacteria and the other with the contour of the detected cell. The test set consists of 87 cases of labelled bright field images unseen by the trained network. The accuracy of the network is calculated over this test set. (total 1513)

### Dataset 2

This dataset is used in training of the semantic segmentation network for segmentation and classification of *Myxococcus xanthus* and *Escherichia coli* from phase contrast images. To separate *M. xanthus* and *E. coli* we designed experiments with a mixed colony of *M. xanthus* and *E. coli*, where *M. xanthus* is tagged with green fluorescence (GFP, DM31, Table S1) and *Escherichia coli* with red fluorescence (mCherry, EC500 -Shaner et al. 2004-Table S1). This dataset is used later for semantic segmentation tasks to separate *M. xanthus* and *E. coli* from phase contrast images (see below).

### MiSiC, Shape-index map based segmentation

The shape index (SI) map of an image, x, calculated over a scale σ, is defined as 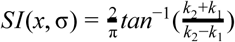, where *k*_1_, *k*_2_ (with *k*_1_ > *k*_2_) are the eigenvalues of the Hessian of the image, *x*, calculated over a scale σ^7^. SIM remains within the range [–1, 1] and preserves the MiSiC shape information while being independent of the intensity values of the original image. Using the Dataset 1, we pre-process the input images X’ to generate a train set: X_train_ of size 1500 × 256 × 256 × 3. Each channel in X_train_ is the shape-index map calculated at scales [1,1.5,2]. An instance of the U-net is trained over this data set to produce a network able to map data represented by X_train_ → y_train_. The network learns to reject the noise in the shape-index map and produces masks and boundaries of the cell like structures in the shape-index map. The trained network was tested over 176 cases of labelled bright-field images, that were previously unseen by the network leading to a segmentation accuracy of 0.76 computed with Jaccard coefficient ^18^.

### Preprocessing

Preprocessing the input image to enhance the edge contrast and homogenising intensities helps in obtaining a good segmentation via MiSiC. Some of the preprocessing that gave good results are gamma correction for homogenising, unsharp masking for sharpening the image and sometimes a gaussian of laplace of the image that removes intensity variations in the entire image and keeps edge-like features.

### Parameters: Scale and noise variance

The dataset 1 used to train the MiSiC contains cells with a width in the range of 8-10 pixels.Thus, to obtain a satisfactory segmentation, the input image must be scaled so that the average bacteria width is around 10 pixels. However, the scaling often modifies the original image leading to a smoother shape index map. Since, MiSiC has basically learned to distinguish between smooth curvatures with well-defined boundaries from noisy background. A smooth image without inherent noise leads to a lot of false positive segmentations. Therefore, counterintuitively, synthetic noise must be added to the scaled or original image for a proper segmentation. It must be kept in mind that the noise variance should not reduce the contrast of the edges in the original image while it should be enough to discard spurious detections. Gaussian noise of a constant variance may be added to the entire image or alternatively, the variance could be a function of the edges in the input image.

### Semantic segmentation: *Myxococcus xanthus* and *Escherichia coli*

The fluorescence images from Dataset 2 are processed with a gamma adjustment and segmented using MiSiC to produce clean masks and contours of two classes, namely, *Myxococcus xanthus* and *Escherichia coli*. Thus, y in dataset 2 contains two channels corresponding to a mask of each class. Another U-NET is then trained on this data to segment a single channel phase-contrast image into an image containing semantic classification of each species. The probability map for each label is color coded (blue = *E. coli*, orange = *M. xanthus*) such that each pixel has a probability value top be part of a given class. In rare instances, bi-color objects are obtained because in these cases the prediction is not homogeneous inside the predicted objects. These objects were filtered for the morphometric analysis showed in Figure 4c.

### Image analysis and statistics

#### Comparison between MiSiC and semi-manual counts

In Figure S1, a 4×4 mosaic of *Escherichia coli* phase contrast image at 100x objective with an approximate surface of 80.10^3^ µm^2^ was used for comparison. In order to semi-manually count the number of cells, the phase contrast image was analysed with FIJI ^19^ with the following commands:

1. FFT Band-pass filter (0 - 40 pixels)
2. Autothreshold
3. Morphomath opening
4. Median filter
5. Watershed (BioVoxxel plugin) [http://www.biovoxxel.de/development/]
6. Size filter (> 40 pixels)
7. Ultimate erosion points.

The points obtained are superimposed to the phase contrast image and manually checked and corrected (Figure S1 A).

The MiSiC spatial density and threshold mask density were derived directly from the binary masks produced for each method and transformed by ultimate erosion of the points.

Spatial density was calculated with the density plugin for FIJI write by Thomas Bourdier [https://imagejdocu.tudor.lu/plugin/analysis/density_2d_3d/start]) from 3D suite^20^, for each pixel x,y:

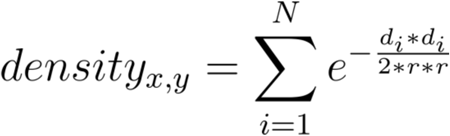

where N = Number of closest neighbours (objects) to compute from each pixel (we used N=20), d_i_ = distance to the closest point, r = the radius of expansion from the spot centre (we used 50 pixels). Units of density are the mean number of objects within a circle with r radius.

The spatial correlation maps (Figure S1) and the correlation graph (Figure 2C) were calculated with Image CorrelationJ (http://www.gcsca.net/IJ/ImageCorrelationJ.html).

#### Analysis of MiSiC performance in the presence of noise and comparison with Supersegger

To illustrate MiSiC performances in the presence of noise and in comparison with SuperSegger^3^, datasets consisting of 141 *E. coli* microcolony images were retrieved from the SuperSegger website. These images were analysed with the provided parameters with SuperSegger and with the following parameters with MiSiC: Cell width = 9, Scaling factor = Auto, Noise =0.001, Unsharp=0.6 and Gamma=0.1. To assess the segmentation robustness to noise for each program, datasets were normalized by the maximum intensity value recorded in the first frame of the dataset and Gaussian noise was added with varying variance. The resulting datasets were then analysed with the initial parameters used to compute reference segmented images except for MiSiC where the noise parameter was set to 0. The relative performance of each program was then evaluated by computing Dice indices^8^.

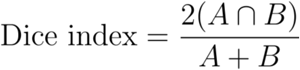

#### Morphological analyses

Classic morphological features: Area, Perimeter, Bounding Box (Width, Height), Circularity, Feret diameter, minimum Feret diameter, MajorAxis (ellipse), MinorAxis (ellipse), (n) number of objects.

Special calculated morphological features: Solidity = Area / Convex Area; AR = MajorAxis / MinorAxis; Extend = Area / (Width*Height)

Morphology index for Figure 2c was calculated by:

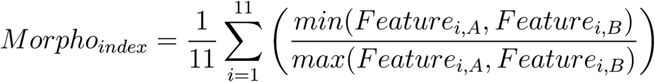

with A = predicted mask, B = manual ground truth. Features = [Area, Perimeter, Circularity, Major, Minor, Feret, minFeret, AR, Solidity, Extend, n_objects_]

#### Demograph construction

To construct the plot shown in Figure 3, the cells bodies were obtained with MiSiC segmentation and the binary mask was analyzed with the MicrobeJ software^1^, with a cell model set to parameters (area > 0.5 µm^2^, Circularity < 0.8, “poles” = 2) to filter all remaining segmented objects that do not correspond to cells. The localization of the centroid (cell middle) and length of longitudinal axis was then determined for each cell under MicrobeJ. The fluorescent clusters were detected with a local maxima filter and their position relative to the middle of the cell was plotted along the axis with a positive sign. Negative sign clusters are therefore the manifestation of rare cells with bi-polar foci. The fluorescent clusters are plotted as dots with a color scale based on spatial density.

#### Cell division detection

In Figure 3e, clusters of fluorescent protein FtsZ-NG were used as cell division markers. The clusters were detected by local maxima detection (scikits-image.peak_local_max(image = fluorescence image, label = MiSiC mask, num_peaks=1))

#### Calculation of cell division ratios

The cell division ratio in Figure 3i was calculated using the spatial 2D density derived from (i) the mask of the total fluorescent cell population across the entire image and (ii), the mask of the total number of fluorescent maxima (reflecting dividing cells) across the entire image. Spatial densities were calculated with sklearn.neighbors.KernelDensity(), with a bandwidth of 1% of the image width. The proportion of dividing cells was obtained by dividing the spatial density maps: density of fluorescent maxima/density of total cells.

## Supplementary information

Supplementary Figure S1. MiSiC can segment *E. coli* micro-colonies and is robust to noise

Supplementary Figure S2. MiSiC segmentation of multiple bacterial species.

Supplementary Figure S3: BactoHubble

Supplementary Table T1: Bacterial strains in this study

Supplementary Table T2: Primers

Supplementary Table T3: Plasmids

### Acknowledgements

We would like to thank Anke Treuner-Lange in Lotte Sogaard-Andersen lab for sharing the plasmid pKA32 with us (Treuner-Lange et al., Mol Micro 2013). MiSiC robustness was tested on microscope images showing various bacterial strains that were kindly shared with us. For that, we would like to thank Anne Galinier and Thierry Doan (*Bacillus subtilis*), Christophe Bordi (*Pseudomonas aeruginosa*), Aretha Fiebig (*Caulobacter crescentus*), Corinne Aubert (*Desulfovibrio vulgaris*) and Romain Mercier (SgmX-GFP images of *Myxococcus xanthus*). We also thank Baptiste Piguet-Ruinet and Célia Jonas for constructing the pDM14 plasmid during their internship. SP, LE and TM are funded by CNRS within a 80-Prime initiative. EM is funded by an AMIDEX-PhD program from Aix-Marseille University. MN, JBF, AL and SR are funded by two ANR grants IBM (ANR-14-CE09-0025-01) and HiResBacs (ANR-15-CE11-0023).

## Code availability

A MiSiC pip installable python package is available at https://github.com/pswapnesh/MiSiC

