## Supplementary material for "MiSiC, a general deep learning-based method for the high-throughput cell segmentation of complex bacterial communities": Figure S1

### Supplementary information

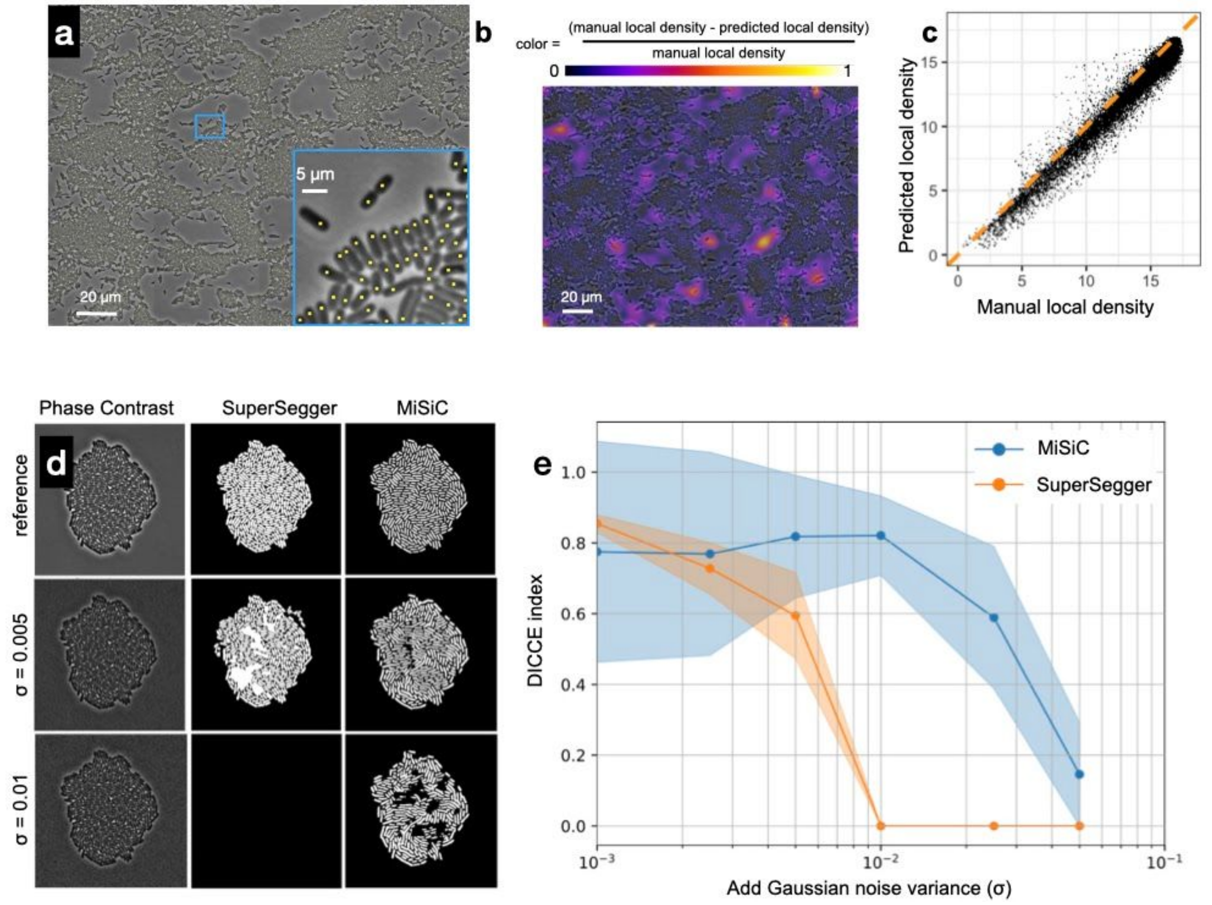


**Figure S1. MiSiC can segment *E. coli* micro-colonies and is robust to noise**

**A-C**: Comparison of MiSiC segmentation with semi-manual counts

(A) 10951 *E. coli* were semi-manually detected in a mosaic image (4x4 fields, see methods) and are shown as yellow dots.

(B) local density subtraction map.The local density maps of the MiSiC mask and semi-counts were computed using in each case the centroid of detected cells. In the subtraction map, the cell density areas have values close to zero ( 0.074) with a standard deviation of 0.07, as expected if there is a good agreement between semi-manual and MiSiC-based density determination.

(C) Correlation plot of the estimated spatial density from semi-manual counts and predicted spatial density obtained through MiSiC. A strong linear correlation over the entire range of spatial densities with a slope of 1.00 with R2 = 0.96 is obtained. Yellow dashed line marks the expected perfect correlation (for an equation y=x).

**D-E:** Robustness of MiSiC to noise and comparison with SuperSegger.

(D) Resulting images with added noise and corresponding segmentation with SuperSegger and MiSiC. To evaluate each program's robustness to noise, an *E. coli* dataset was processed with SuperSegger and MiSiC in the presence of increasing amounts of Gaussian noise added to the images while keeping segmentation parameters constant (Methods).

(E) Relative performance of SuperSegger and MiSiC to increasing noise. For each amount of noise the complete datasets (141 images) were processed and the Dice index (Method) was calculated with respect to the segmentation results (ie panel D). Each dot on the lines represents the mean Dice value while the shaded error bars represent its standard deviation across the dataset. Note that while Superseger and MiSiC perform equally well at low noise (on a Supersegger optimized dataset, Methods), MiSiC remains robust in the presence of noise, while the efficiency of SuperSegger drops very rapidly.
