## Supplementary material for "MiSiC, a general deep learning-based method for the high-throughput cell segmentation of complex bacterial communities": Figure S2

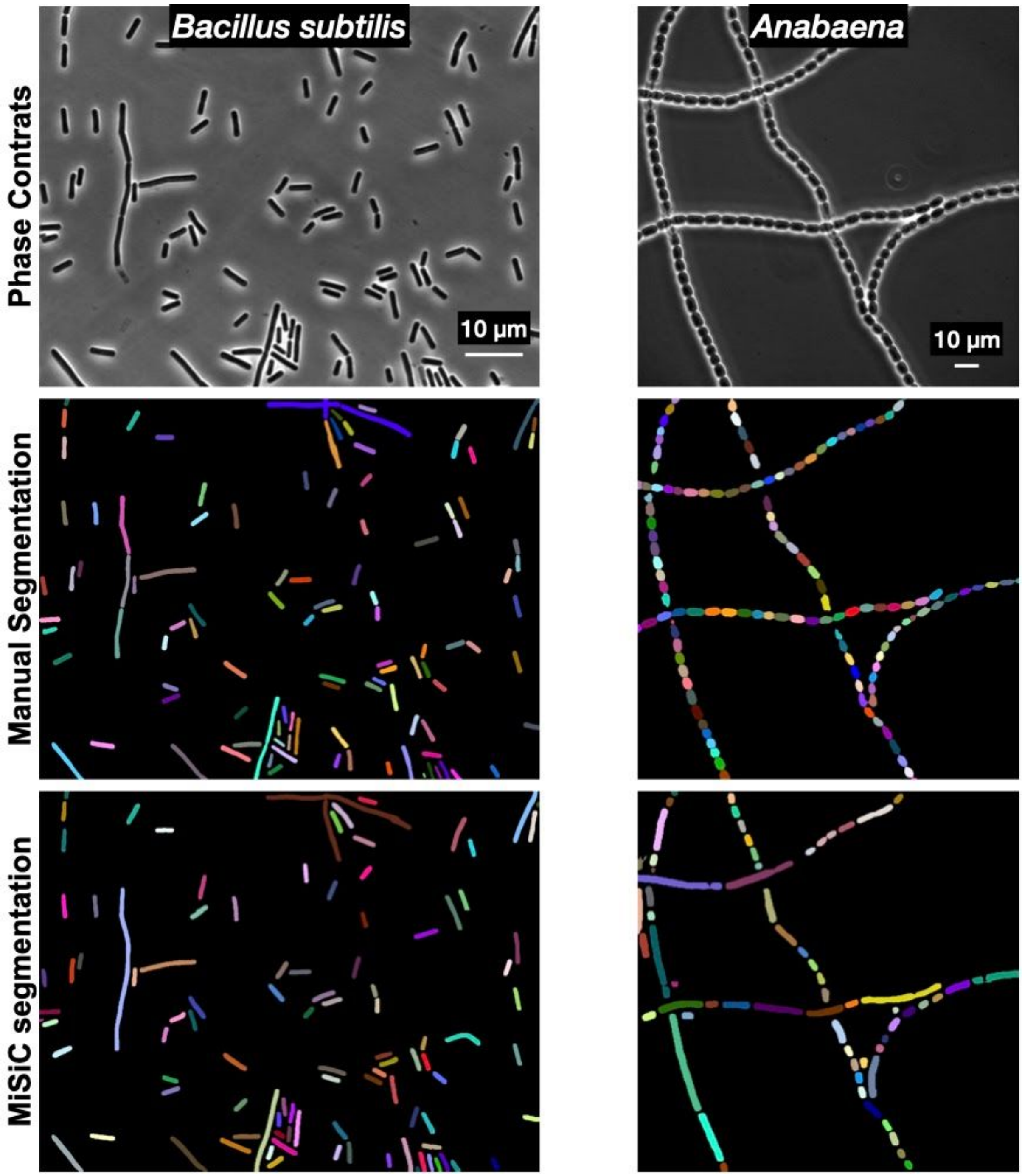


**Figure S2: MiSiC segmentation of multiple bacterial species.** Comparison of manual segmentation and MiSiC segmentation for a typical rod-shaped bacterium *Bacillus subtilis* and a filamentous cyanobacterium, *Anabaena sp*. Note that in both cases, MiSiC errors appear for the segmentation of filaments with multiple septa, structures which are difficult to capture in the SIM image.
