## Supplementary material for "MiSiC, a general deep learning-based method for the high-throughput cell segmentation of complex bacterial communities": Figure S3

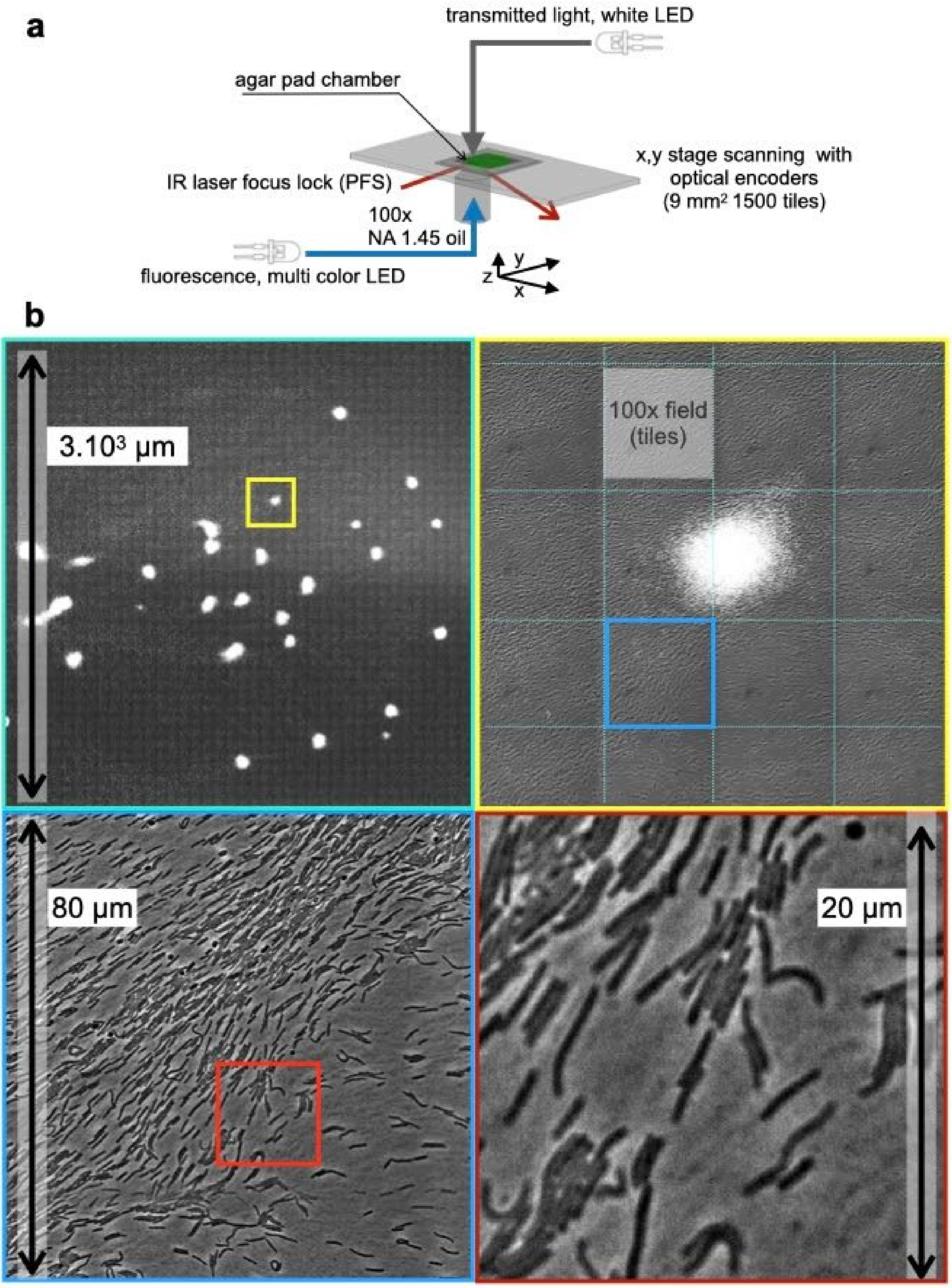


**Figure S3: Bacto-Hubble captures millimeter size images of bacterial communities with single cell resolution.**

**a)** Bacto-hubble microscope set-up. The inverted microscope and the sample holder have been optimized to increase the speed of tile acquisition and the stability of focus. The key point is to replace as many moving parts as possible, such as shutters, and replace them with diode sources. The agar pad is blocked by a flat coverslip that, combined with the continuous focus control (in our case a Nikon PFS system), makes it possible to maintain the focus over millimeter and up to centimeter distances. The motorized stage is equipped with optical encoders, allowing precise x, y movements across the specimen.

**b)** Bacto-Hubble image of an entire predator-prey colony. The area shows an *M. xanthus* community 96 hours after it started invading an *E. coli* prey colony. At this stage, the prey has been entirely consumed. Following growth and starvation, *M. xanthus* cells aggregate (phase-bright dots in the image, top panels) and form fruiting bodies. The entire image corresponds to the assembly of 40x40 tiles for a total surface of 9 $mm^2$. Top right panel is a close up of the region framed in light blue in the top left panel. Bottom left panel is a close up of the region framed in yellow in the top right panel. Bottom right panel is a close up of the region framed in red in the bottom left panel showing individual *M. xanthus* cells.
