## Supplementary material for "MiSiC, a general deep learning-based method for the high-throughput cell segmentation of complex bacterial communities": Table T1

**Table T1 Bacterial strains used in this study**

| Strain name | Strain/Genotype | Origin |
|---|---|---|
| *Myxococcus xanthus* | | |
| DZ2 | | Laboratory collection |
| DM14 | DZ2 *attmx8 ftsZ*-NG | This work |
| DM31 | DZ2 p3068-sfGFP | This work |
| *Escherichia coli* | | |
| MG1655 | | Laboratory collection |
| EC500 | TOP 10 pFPV-mCherry (pGG2-rpsM-mCherry) | Laboratory collection |
| *Pseudomonas aeruginosa* | Wild type strain PAK | Laboratory collection |
| *Bacillus subtilis* | Prototrophic wild-type strain, 168CA *trpC*+ | Laboratory collection |
| *Bacillus subtilis* | Wild type, Strain 168 | Laboratory collection |
| *Caulobacter crescentus* | NA1000-*hfsA*+ | Marks M.E et al. J. Bact. 2010[15] |
| *Anabaena nostoc* | PCC 7120 | Pasteur Institute cyanobacterial collection |
| *Desulfovibrio vulgaris* | Wild type strain | Laboratory collection |
