## Supplementary material for "MiSiC, a general deep learning-based method for the high-throughput cell segmentation of complex bacterial communities": Table T2

**Table T2. Primers**

| Primer name | Sequence | Use | Construct |
|---|---|---|---|
| oDM1 | CTC-TAG-AAC-TAG-TGG-ATC-CGA-ACA-ACC-GCC-GCG-TGG-GG | cloning of *ftsZ* | pDM6 |
| oDM2 | GTA-TTT-CAC-ACC-GCA-TAT-GTT-ACG-GCA-GTT-CCG-TCT-GGC | cloning of *ftsZ* | pDM6 |
| oDM16 | cct-gca-ggt-cga-ctc-tag-atc-act-tat-aga-gtt-cat-cc | cloning of Neon-Green | pDM6 |
| oDM17 | gac-gga-act-gcc-ggg-tac-cgg-tac-cgg-gcc-ccc-cct-c | cloning of Neon-Green | pDM6 |
| oDM53 | GTT-CTT-CAC-CTT-TAG-ACA-TTG-ACA-CTC-CTC-AAA-AAT-AAA-TGG-A | cloning of *MXAN_3068* upstream region | pDM14 |
| oDM54 | GGG-GAT-CCG-GGC-GAA-CGG-GAA-TTC-TA | cloning of *MXAN_3068* upstream region (p3038) | pDM14 |
| oDM61 | CCG-CAT-ATG-TTA-TTT-GTA-GAG-CTC-ATC-C | cloning of sfGFP downstream of p3068 | pDM14 |
| oDM62 | TTT-TGA-GGA-GTG-TCA-ATG-TCT-AAA-GGT-GAA-GAA-C | cloning of sfGFP downstream of p3068 | pDM14 |
