## Supplementary material for "MiSiC, a general deep learning-based method for the high-throughput cell segmentation of complex bacterial communities": Table T3

**Table T3. Plasmids**

| Plasmid name | Backbone | Genotype |
| --- | --- | --- |
| pDM6 | pKA32 (Treuner-Lange, Mol. Micro 2013) | Pnat-*ftsZ*-linkerNeonGreen |
| pDM14 | pSWU19 | pSWU19_*p3068*-sfGFP |
